# Heat shock factor 2 positively regulates oncogenic herpesvirus gene expression by remodeling the chromatin landscape

**DOI:** 10.1101/2024.11.22.624798

**Authors:** Lorenza Cutrone, Hedvig Djupenström, Jasmin Peltonen, Elena Martinez Klimova, Simona Corso, Silvia Giordano, Lea Sistonen, Silvia Gramolelli

## Abstract

Human gamma-herpesviruses, Kaposi’s sarcoma herpesvirus (KSHV) and Epstein-Barr virus (EBV) are causally associated to a wide range of cancers. While the default infection program for these viruses is latent, sporadic lytic reactivation supports virus dissemination and oncogenesis. Despite its relevance, the repertoire of host factors governing the transition from latent to lytic phase is not yet complete, leaving much of this complex process unresolved. Here, we show that heat shock factor 2 (HSF2), a transcription factor involved in regulation of acute stress responses and specific cell differentiation processes, promotes gamma-herpesvirus lytic gene expression. In lymphatic endothelial cells infected with KSHV and gastric cancer cells positive for EBV, ectopic HSF2 enhances the expression of lytic genes, while knocking down HSF2 significantly decreases their expression. Mechanistically, HSF2 overexpression results in decreased levels of repressive chromatin histone marks, at the promoters of the master regulators of the oncogenic lytic cascade, KSHV ORF50 and EBV BZLF1. Our results demonstrate that endogenous HSF2 binds to the ORF50 promoter in latent cells and sustains a transcriptionally permissive state. In contrast, in lytic cells, HSF2 occupancy at the ORF50 promoter is lost in conjunction with its proteasomal degradation. These findings establish HSF2 as a conserved regulator of gamma-herpesvirus lytic gene expression in latency and offer new functional insights on the cellular role of HSF2 at target promoters.

## Introduction

The gamma-herpesviruses Kaposi’s sarcoma associated herpesvirus (KSHV) and Epstein-Barr virus (EBV) are human oncogenic herpesviruses, recognized by the World Health Organization (WHO) as class I carcinogens. Together they account for approximately 2-3% of all cancer cases worldwide (www.globocan.com). KSHV seroprevalence varies globally, with higher rates in regions like sub-Saharan Africa and the Mediterranean basin. KSHV is causally associated to primary effusion lymphoma, multicentric Castleman’s disease and Kaposi’s sarcoma (KS)(1,2). EBV is a ubiquitous virus, infecting 95% of the human population and it was the first pathogen etiologically associated with human malignancies (3). EBV infection is linked to various types of cancers, including endemic Burkitt’s lymphoma, nasopharyngeal carcinoma (NPC), and EBV-associated gastric cancer (EBVaGC), a recently identified molecular subtype of gastric cancer (3–7).

While the default infection program for gamma-herpesviruses is latency, sporadic lytic reactivation episodes are necessary for viral dissemination (8,9). During latency, the viral genomes persist in the nucleus of the infected cells as circular episomes which are epigenetically silenced. The gene expression is restricted to a handful of latent genes that are fundamental for evading the immune surveillance and ensure host cell survival (10,11). In contrast, during the lytic phase, all viral genes are expressed according to a well-orchestrated temporal cascade. The process initiates with the expression of early lytic genes that promote viral replication, followed by the late lytic genes needed for the assembly and release of infectious viral progeny. (8,9,12,13). Clinical observations highlight the important contribution of lytic reactivation in tumorigenesis, as KS patients undergoing treatment with inhibitors of the lytic replication experience tumor regression (14). Essential for initiation and completion the lytic cycle are the key transcription factors RTA and Zebra 1, encoded by KSHV ORF50 and EBV BZLF1, respectively (12). Although it is known that the expression of these viral factors is necessary and sufficient to induce the full lytic cascade, the molecular events governing their regulation remain still obscure (12,15).

Environmental stimuli, like hypoxia and oxidative stress, have been implicated in controlling the switch from latency to productive lytic replication of several herpesviruses (12,16). These stressors stimulate, in turn, the expression of heat shock proteins, including HSP70 and HSP90, which function as molecular chaperones and are often hijacked by the viral replication machineries (17,18). HSPs are host factors required for productive viral replication, where large amounts of essential, structural viral proteins need to be appropriately folded (18). Additionally, both HSP70 and HSP90 are associated with the newly produced EBV and KSHV viral particles (19). HSP70 has been also shown to relocalize from the cytoplasm to the nucleus, where it forms dynamic structures adjacent to the viral replication and transcription compartments during KSHV lytic phase (20).

In response to stress, the expression of genes encoding HSP70 and HSP90 is induced by the specific members of the heat shock factor (HSF) family, *i.e.* HSF1 and HSF2. HSF1 is the primary mediator of the acute stress responses, whereas HSF2 supports HSF1-mediated stress-related functions (21,22). Upon acute stress, HSF1 undergoes extensive post-translational modifications, causing its release from the complex with HSPs, and forms either homotrimers or heterotrimers with HSF2 (23,24). These HSF complexes drive the transcription of downstream genes by binding to specific *cis*-acting genomic motifs known as the heat shock response elements (HSEs). Depending on the biological context, HSFs can function both together and independently from one another. In stress response, for example, HSF1 and HSF2 share common targets, but different subsets of genes regulated exclusively by either HSF1 or HSF2 have also been identified (25,26). In many solid tumors, HSF1 and HSF2 interact and together modulate subsets of genes fundamental for disease progression, whereas in breast cancer HSF2 alone has been shown to promote the abnormal proliferation of cancer cells (27,28). In addition to regulating cellular stress responses and malignant progression, HSF2 has also been implicated in diverse cellular processes ranging from erythroid differentiation to spermatogenesis and corticogenesis (21,24,25,29–31). In the context of herpesviral infections, recent studies have demonstrated that HSF1 activation is required for efficient replication of human herpesvirus 6A and human cytomegalovirus (HCMV) (32,33). Furthermore, in EBV-infected cells, HSF1 is recruited to an HSE-enriched region within the promoter of the EBV latent gene EBNA-1, thereby promoting its expression (34). In this study, we focus on HSF2 whose role in viral infection has remained unexplored.

Here we provide evidence for HSF2 acting as a positive regulator of KSHV and EBV gene expression programs. We found that HSF2 specifically regulates the KSHV ORF50 promoter in an RTA and HSF1-independent manner. Interestingly, HSF2 binds the ORF50 promoter in latent but not lytic cells and undergoes proteasomal degradation upon induction of KHSV lytic cycle. In the presence of ectopically expressed HSF2, the chromatin at the KSHV ORF50 and EBV BZLF1 promoters showed decreased repressive marks, indicative of a more permissive viral transcriptional state during latency. This study expands our understanding of the complex interplay between gamma-herpesviruses and the host cellular machinery.

## Results

### HSF2 expression modulates the KSHV lytic cycle in KSHV-infected lymphatic endothelial cells

To investigate the potential role of HSF2 in KSHV life cycle, we perturbed its expression in KSHV-infected lymphatic endothelial cells (KLEC). KLEC have the unique property to support a spontaneous KHSV lytic expression program. In these cells, lytic expression as well as infectious progeny production and release occur without any trigger. We decided to use this cell model as KLEC are putative cells of origin of KS, thus representing a relevant *in vitro* model for studying the regulatory mechanisms of KSHV gene expression and tumorigenesis (8,35–37).

First, we overexpressed either HSF2 or a GFP control vector by lentiviral transduction in KLEC for 72 h and demonstrated elevated HSF2 protein levels by immunoblot (Fig. 1A). We observed a pronounced increase in KHSV early (K-bZIP) and late (K8.1) lytic protein expression (Fig. 1A) along with a concomitant elevated expression of lytic genes including *ORF57* and *ORF50* (Fig. 1B). HSF2 overexpression also enhanced secretion of viral particles from the infected cells (Fig. 1C). Subsequently, we tested the effect of HSF2 downregulation on KSHV expression and virus release. KLEC were transiently transfected with either a control (scr) siRNA or an siRNA targeting HSF2 for 72 h. Efficient silencing of HSF2 and consequent protein downregulation was demonstrated by immunoblot (Fig. 1D). A significant reduction in KSHV lytic protein levels and gene expression was observed (Fig. 1D, E), along with diminished virus titers (Fig. 1F). Altogether, these findings show that perturbing HSF2 levels influences KSHV gene expression, thereby pointing to HSF2 as a novel positive regulator of KSHV gene expression and viral production in primary KLEC.

**Figure 1.**
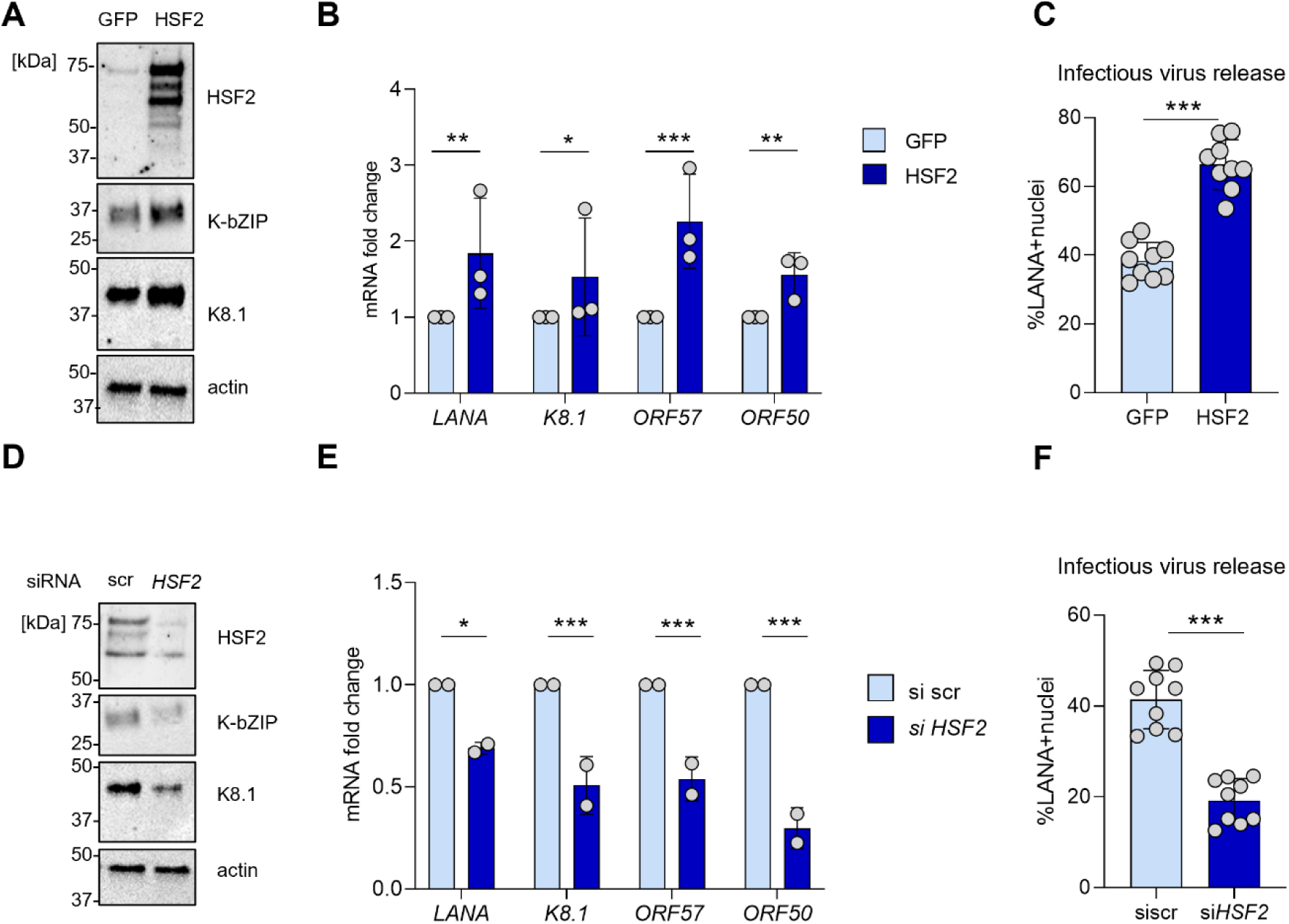
HSF2 expression sustains KSHV infection in KLEC. Primary LEC were infected with KSHV for 14 days. (A, B, C) KLEC were transduced with lentiviruses carrying either a GFP or a HSF2 gene for 72 h prior to analysis. (D, E, F) KLEC were treated with siRNA control (scramble, scr) or targeting HSF2 for 72 h prior to analysis. (A, D) Representative Immunoblot analysis of HSF2 and viral K-bZIP and K8.1 proteins, actin was used as a loading control. Molecular weight in kDa is shown on the left. (B, E) RTqPCR analysis of the indicated transcripts. *Actin* was used as a internal control. Transcript fold changes relative to GFP (B) siscr (E) are shown. Bars indicated average, error bars the standard deviation across three independent experiments, single data points are indicated as grey circles (C, F) the supernatant from KLEC treated as indicated was serially diluted and applied to uninfected U2OS cells for 24h. Cells were stained for LANA, nuclei were counterstained with DAPI and imaged with high throughput microscope. Percentage of LANA positive nuclei are shown. Bars indicated average, error bars the standard deviation across independent experiments, single data points are indicated as grey circles.

### HSF2 regulates the KSHV lytic gene expression during latent and lytic phases

To determine whether HSF2 modulates KSHV gene expression during latency or at the onset of the lytic phase, we opted to use the iSLK.219 cells, a different cell-based system from KLEC. Although representing a physiologically relevant model of KSHV infection, KLEC feature a continuous and asynchronous lytic reactivation. Therefore, in KLEC, the cellular events occurring in latency cannot be distinguished from those characterizing the lytic replication. Instead, iSLK.219 cells sustain a tight latent infection program. These cells have been engineered to harbor an ectopic, doxycycline inducible *ORF50* gene for the expression of viral RTA, necessary and sufficient to induce the lytic cycle. In addition, iSLK.219 cells are infected with a recombinant rKSHV.219 virus which harbors a constitutively expressed GFP gene and an RFP reporter expressed only during lytic cycle, controlled by the early viral lytic polyadenylated nuclear RNA (PAN) promoter (38,39). Thus, iSLK.219 cells represent an optimal model for accurate monitoring of the lytic reactivation which can be efficiently induced by doxycycline treatment.

We used lentiviral transduction to overexpress either HSF2 or a GFP control in latent iSLK.219 cells for 48 h followed by a 24-h treatment with doxycycline. Immunoblot analysis confirmed higher levels of HSF2 upon overexpression and showed an increase in early and late lytic viral proteins, K-bZIP and K8.1, respectively (Fig. 2A). Moreover, we observed more RFP-positive cells (Fig. 2B) when HSF2 was overexpressed. The augmented lytic reactivation was also measured at the transcriptional level, with significantly higher *ORF57* (early lytic) and *K8.1* gene expression in cells with ectopic HSF2 (Fig. 2C). To validate our results, we investigated the effect of transient HSF2 depletion during the lytic phase in iSLK.219 cells. To this aim, we used siRNA targeting HSF2 or a control siRNA (siscr) for transfecting iSLK.219 cells and 48 h later we induced the lytic cycle by doxycycline treatment for 24 h. Immunoblot analysis showed a robust HSF2 downregulation and lower levels of the K-bZIP viral protein (Fig. 2D). Additionally, HSF2 depletion led to a decrease in RFP-positive cells (Fig. 2E) and in viral gene expression (Fig. 2F). These results confirm also in lytically induced iSLK.219 cells that HSF2 is a positive modulator of KSHV gene expression.

**Figure 2.**
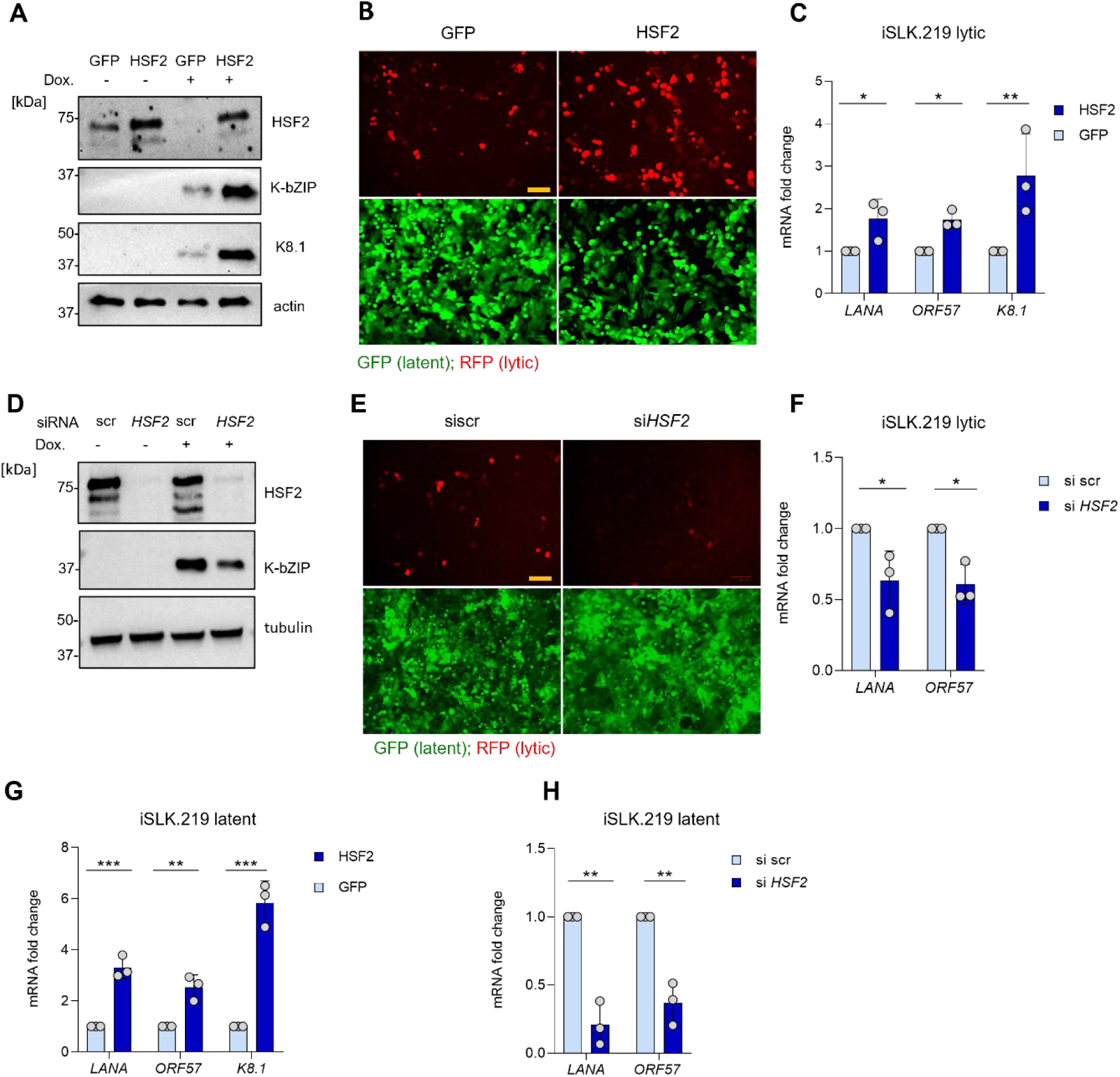
HSF2 regulates lytic gene expression in latency and during the lytic phase. (A) iSLK.219 cells were transduced with GFP or HSF2 encoding lentiviruses for 48 h, where indicated Doxycycline (DOX) treatment was added for 24h. Cell lysate was analysed by immunoblot for HSF2 and for the viral proteins K-bZIP and K8.1. Actin was used as a loading control. Representative blots are shown. Molecular weight in kDa is shown on the left. (B) Fluorescent microscopy images of iSLK.219 cells treated as in (A) and reactivated with Doxycycline for 24 h. Scale bar 100 μm. (C) iSLK.219 cells were treated as in (A) and analyzed by RTqPCR for the indicated viral transcripts. Transcript fold changes relative to GFP are shown. *Actin* was used as an internal control. Bars indicate the average and the error bar the SD across three independent experiments, single data points are shown as grey circles. (D) Representatives immunoblot analysis of iSLK.219 cells transfected with the indicated siRNAs for 48 h, where indicated Doxycycline (Dox) treatment was added for 24h. Cell lysate was analysed by immunoblot for HSF2 and for the viral protein K-bZIP. *Tubulin* was used as a loading control. Molecular weight in kDa is shown on the left. (E) Fluorescent microscopy images of iSLK.219 cells treated as in (D) and reactivated with Doxycycline for 24 h. Scale bar 100 μm. (F) iSLK.219 cells were treated as in (E) and analyzed by RTqPCR for the indicated viral transcripts. Transcript fold changes relative to si scr are shown *Actin* was used as an internal control. Bars indicate the average and the error bard the SD across three independent experiments, single data points are shown as grey circles. (G, H) iSLK.219 cells were transduced with the indicated lentiviruses for 48 h (G) or transfected with the indicated siRNA (H). Transcript fold changes relative to GFP are shown. *Actin* was used as an internal control. Bars indicate the average and the error bard the SD across three independent experiments, single data points are shown as grey circles.

Since the effects of HSF2 modulation on KSHV expression were evident already at 24 h after doxycycline induction, we asked whether HSF2 would influence the levels of lytic viral gene expression prior to the induction of the lytic cycle, *i.e.* in latently infected cells. To answer this question, we measured viral gene expression in latent iSLK.219 cells overexpressing HSF2 for 48 h and observed significantly increased expression of LANA, ORF57 and K8.1 (Fig. 2G). Conversely, silencing HSF2 in latent iSLK.219 cells led to significantly reduced levels of viral transcripts (Fig. 2H). Together, the results reveal that HSF2 enhances KSHV gene expression in latent cells, hence playing a regulatory role in both lytically reactivated iSLK.219 cells and during latency.

### HSF2 regulates the KSHV ORF50 promoter independently of RTA and HSF1

The effect of HSF2 on the KSHV gene expression was demonstrated at the transcriptional level. Therefore, we reasoned that HSF2 could enhance KSHV gene expression by regulating viral promoter activities. We decided to address this hypothesis by performing luciferase reporter assays. We selected a set of plasmids harboring the reporter gene downstream of individual viral promoters which were co-transfected with either HSF2 or GFP control expressing vectors. Transfection efficiency and successful HSF2 overexpression were assessed by immunoblot against HSF2 (representative blots are shown in Suppl. Fig. 1 A-D). In these experiments, we could not observe any significant variation in the activity K8.1 or ORF45 promoters in the presence of ectopically expressed HSF2 (Fig. 3A). Similarly, HSF2 overexpression had no significant impact on the activity of either the 7XTR promoter, containing 7 copies of KSHV latent origin of viral replication, or on the KSHV origin of lytic replication, ORILYT (Fig. 3B). However, we detected a strong increase in the activity of the ORF50 promoter (about 60-fold over the GFP control) in conjunction with HSF2 overexpression (Fig. 3C).

**Figure 3.**
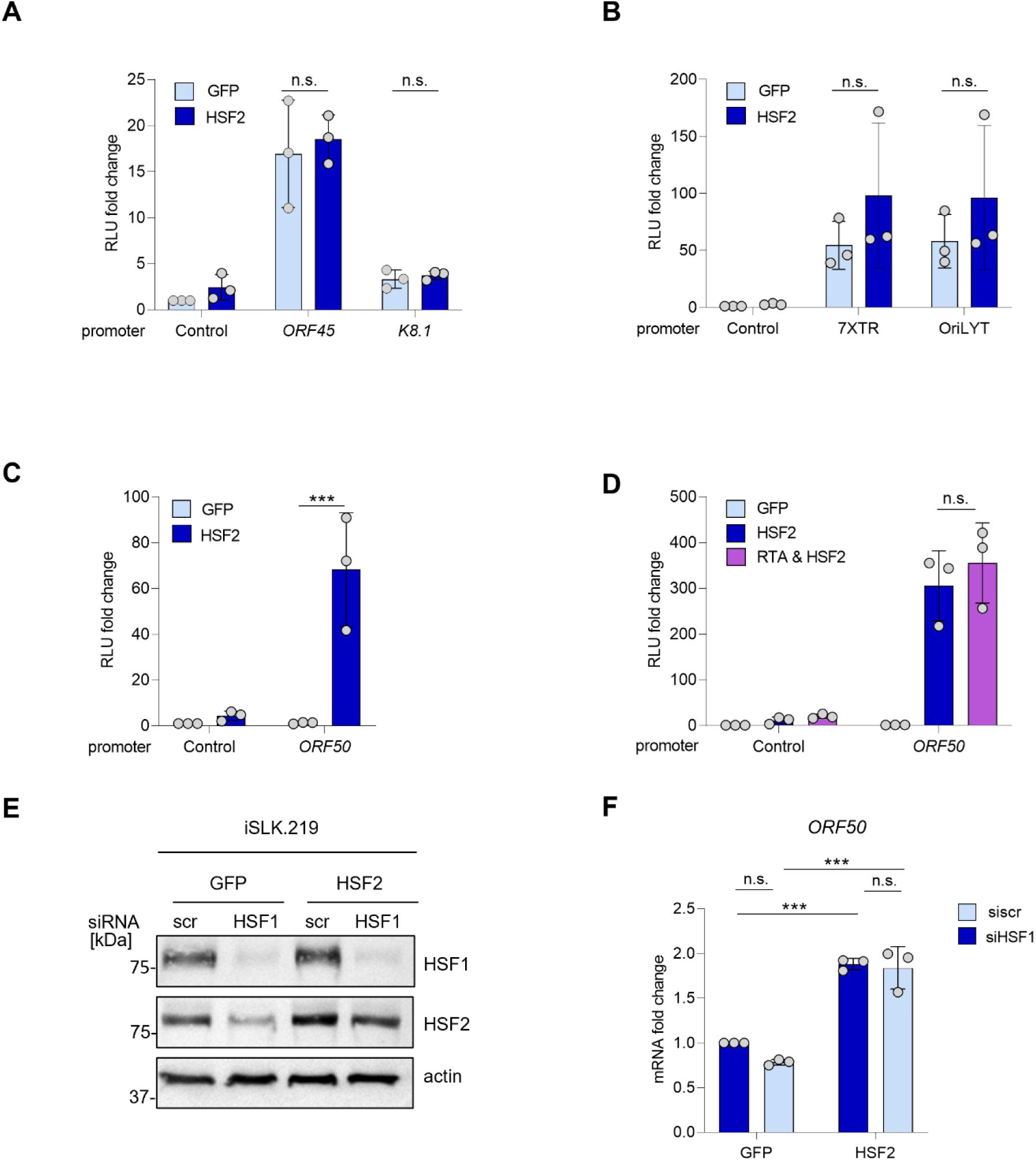
HSF2 regulates ORF50 expression in an RTA independent manner. (A-D) Luciferase reporter assay in HEK293FT cells co transfected in triplicate with a plasmid harboring the luciferase reporter gene downstream of a control basic promoter (control) or the indicated viral promoters (A, C, D) or viral origins of replication (B), along with HSF2 or GFP control expressing plasmids. In (D) cells were also transfected with an RTA-encoding plasmid. Luciferase activity was measured 48h after transfection and indicated as relative luciferase units (RLU) fold change compared to the control -GFP sample. The experiments were repeated three independent times, each datapoint (grey circle) represents the average of the three technical replicates, bars represent the average across three biological replicates, and the error bars the SD across three independent biological replicates. (E) Representatives immunoblot analysis for HSF1 and HSF2 in iSLK.219 cells overexpressing GFP or HSF2 and transfected with either control (scr) siRNA or siRNA targeting HSF1 for 48 hours. *Actin* was used as loading control. Molecular weight in kDa is shown on the left. (F) RTqPCR analysis of cells treated as in (E) for *ORF50* gene, *Actin* was used as loading control. Transcript fold changes relative to GFP, siscr-treated sample are shown. *Actin* was used as an internal control. Bars indicate the average and the error bar the SD across three independent experiments, single data points are shown as grey circles.

ORF50 codes for RTA, the master transcriptional regulator of the KSHV lytic cycle. Beside other downstream lytic promoters, RTA also binds to its promoter and enhances its activity in a positive feedback loop (40). This is fundamental to enhance the efficiency of the lytic reactivation thus ensuring its progression and successful completion (15,40,41). Several cellular transcription factors have been implicated in assisting RTA to further enhance this positive feedback loop. Examples include cellular PROX1, which is expressed in lymphatic endothelial cells and functions as a key contributor to the spontaneously lytic KSHV expression program observed in the lymphatic cellular environment (37). To test whether HSF2 would synergize with RTA, we co-expressed both proteins together with the luciferase ORF50 reporter. KSHV RTA significantly enhanced the activity of ORF50 promoter, in line with previously published results (42). However, the presence of HSF2 did not increase this effect (Fig 3D). Overall, from these results we concluded that HSF2 promotes exclusively and specifically the expression of ORF50 by enhancing its promoter activity in the absence of RTA protein expression.

HSF2 is a transcription factor that can act either directly on its target genes or together with HSF1 (21,26). During acute stress the interplay between HSFs sustains an optimal cellular response, whereas in other physiological and pathological processes such as erythroid differentiation and breast cancer progression, HSF2 operates independently from HSF1 (21,28,30). To address whether HSF2 would act in concert with HSF1 in regulating the KSHV life cycle, we transiently silenced HSF1 for 48 h in latent iSLK.219 cells expressing either ectopic HSF2 or GFP control. Immunoblot analysis showed that upon robust silencing of HSF1, HSF2 levels were also diminished (Fig. 3E), which is in line with previous reports documenting lower HSF2 levels in cells devoid of HSF1 (22,23). In contrast, when ectopically reintroduced, HSF2 levels were not as dramatically affected by HSF1 depletion. In these cells, ORF50 expression was upregulated upon HSF2 overexpression and depletion of HSF1 did not change this increase (Fig. 3F). In U2OS osteosarcoma cells stably infected with KSHV, where a less robust HSF1 depletion was obtained, we confirmed that HSF2 overexpression increased ORF50 levels regardless of HSF1 downregulation. (Suppl. Fig. 1 E, F).

Collectively, these results reveal that in the context of the KSHV life cycle, HSF2 enhances the activity of the ORF50 promoter independently of both RTA and HSF1. The ability of HSF2 to affect ORF50 expression in the absence of RTA supports our observation that HSF2 can modulate lytic genes already in latent cells, when RTA is absent. Importantly, we also show that during latent KSHV infection HSF2 levels are controlled by HSF1, similarly to what has been shown for uninfected cells. Furthermore, our results demonstrated that HSF2 is capable of regulating ORF50 expression in an HSF1-independent manner. This undergoes a separate mechanism that diverges from the previously reported roles of HSF1 in controlling other herpesviral life cycles and gene expression (32–34).

### HSF2 binds the ORF50 promoter and modifies the local epigenetic landscape

HSF2 protein recognizes classic heat shock elements (HSEs) on the promoter of its target genes (43). The HSEs contains a variable number of the pentameric sequence nGAAn arranged in alternate orientation (44). We used the JASPAR database to screen the ORF50 promoter for presence of this motif. We identified a putative HSE displaying a stretch of three nGAAn pentamers located in the region from 546 to 559 nucleotides upstream of the ORF50 transcriptional start site (TSS) (the position of the HSE motif within the ORF50 promoter is highlighted in yellow on the schematics in Fig. 4A, and its nucleotide sequence is shown below along with the HSE sequence logo) To experimentally test the possible binding of HSF2 to the ORF50 promoter, we performed a chromatin immunoprecipitation (ChIP) assay followed by RTqPCR (ChIP-PCR). We used a polyclonal HSF2 antibody (22) to precipitate HSF2 and amplified a region covering about 800 base pairs (bp) upstream of the ORF50 TSS (Fig. 4A). Endogenous HSF2 was bound to the ORF50 promoter in latency (Fig. 4B). To our surprise, in iSLK.219 cells reactivated for 24 h, no HSF2 binding was detected (Fig. 4C), indicating a possible displacement of HSF2 during the lytic replication cycle. As control, we tested also other regions of the KSHV genome, one localized in proximity of the ORF73 gene, coding for the key latent protein LANA and the other localized within the late lytic K8.1 promoter. In these regions, we were not able to detect any HSF2 binding either in latent or in lytic cells (Supplementary Fig.2 A, B).

**Figure 4.**
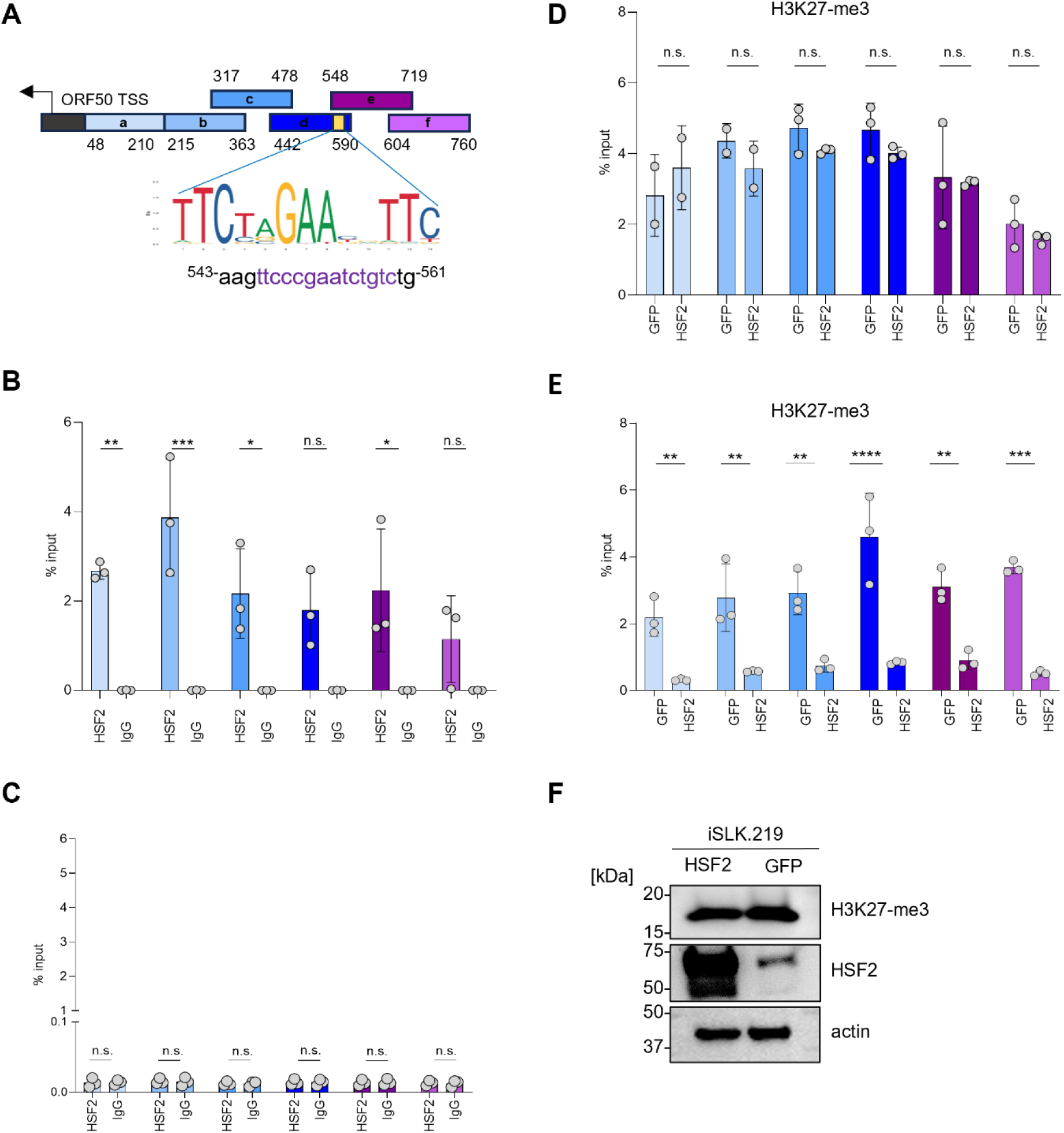
HSF2 binds the ORF50 promoter and remodels the local chromatin landscape. (A) Schematics of the ORF50 promoter region, transcription start site (TSS, black arrow) and HSF2 binding site (yellow box) are shown. The different genomic regions amplified by RTqPCR are depicted as blue and purple segments, the nucleotide positions relative to the TSS are labeled. The DNA motif and the nucleotide sequence identified as HSF2 consensus sequence, along with the HSE sequence logo are shown below. (B, C) ChIP-qPCR analysis of HSF2 binding to the ORF50 promoter regions expressed as % of input in KSHV infected latent(B) and reactivated with doxycycline for 24 h (C). (D, E) ChIP PCR analysis of the indicated histone H3 modifications in latent iSLK.219 cells overexpressing HSF2 or GFP. The bars are colored according to the schematic in (A). Bars represent the average and error bars the SD across three independent experiments, data points are indicated as grey circles. (F) Representatives immunoblot analysis of latent iSLK.219 cells treated as in (D, E) for H3K27-me3 histone, actin was used as a loading control. Molecular weight in kDa is shown on the left.

Epigenetic profiling of the KSHV genome has revealed that, during latency, the KSHV ORF50 promoter bears concomitant activating and repressive histone marks in its chromatin structure characterized by the deposition of tri-methyl groups on either lysine 4 or 27 of the histone H3 (H3K4-me3 and H3K27-me3, respectively) (45). This bivalent or poised state is commonly found in pluripotent stem cells, at the promoters of genes regulating development (46). Under unstimulated conditions, poised gene promoters remain transcriptionally repressed. However, upon receiving appropriate stimuli and concurrent removal of repressive marks, these promoters can rapidly become active thereby leading to gene expression (46,47). Similarly, in the latent phase, the KSHV ORF50 promoter, which bears chromatin markers indicative of a poised state, remains silenced. Upon activation of the lytic cycle, the ORF50 promoter undergoes rapid activation with the removal of the repressive H3K27-me3 chromatin mark (11,45). We then asked whether HSF2 would affect the bivalent chromatin state of the ORF50 promoter. For this, we performed a ChIP-PCR analysis in latently infected iSLK.219 cells overexpressing either HSF2 or GFP control lentiviruses. We used antibodies against H3K27-me3 and H3K4-me3, epigenetic marks for repressive and active chromatin, respectively. While the levels of activating H3K4-me3 were unchanged upon HSF2 overexpression across the ORF50 promoter (Fig. 4D), the H3K27-me3 repressive chromatin mark was significantly diminished (Fig. 4E). As control, we tested the levels of H3K4-me3 and H3K27-me3 in the KSHV genomic regions where HSF2 was not binding, and no change in the chromatin status was detected in cells overexpressing HSF2 (Suppl. Fig 2 C, D). Furthermore, the total levels of H3K27-me3 modifications upon HSF2 ectopic expression remained unchanged as demonstrated by immunoblot (Fig. 4F). This data indicates that the changes in the levels of H3K27-me3 are localized to specific viral genomic loci rather than being broadly distributed.

In conclusion, these results show that HSF2 associates with the ORF50 promoter iSLK.219 cells during latency but not in the lytic phase. In latent cells HSF2 overexpression shifts the bivalent chromatin state of the ORF50 promoter by reducing locally H3K27-me3 chromatin mark, thus priming the viral genome to transition from latency to lytic reactivation.

### HSF2 protein is degraded during the lytic reactivation

The remarkable loss of HSF2 occupancy to the ORF50 promoter 24 h after induction of the lytic cycle prompted us to investigate whether this was due to downregulation of HSF2 levels or to the mislocalization of the protein. To carefully address these hypotheses, we performed both immunofluorescence staining and immunoblotting in iSLK.219 cells that were either latent or reactivated for 24 h. Immunofluorescence revealed a nuclear localization of HSF2, in line with its transcription factor activity. When comparing the integrated nuclear intensity of the immunofluorescence signal, we measured a significant reduction of HSF2 intensity in the lytic cell population, compared to the latently infected cells (Fig. 5A, B). We next monitored HSF2 levels by immunoblot during lytic reactivation in iSLK.219 during a 48-h time-course (Fig. 5C). We observed a slight increase in HSF2 levels as soon as 6 h after the induction of the lytic cycle, followed by an initial decrease at 24 h and a more dramatic downregulation of HSF2 at 48 h after reactivation. Together these results suggest that HSF2 protein is downregulated rather than mislocalized during KHSV lytic cycle.

**Figure 5.**
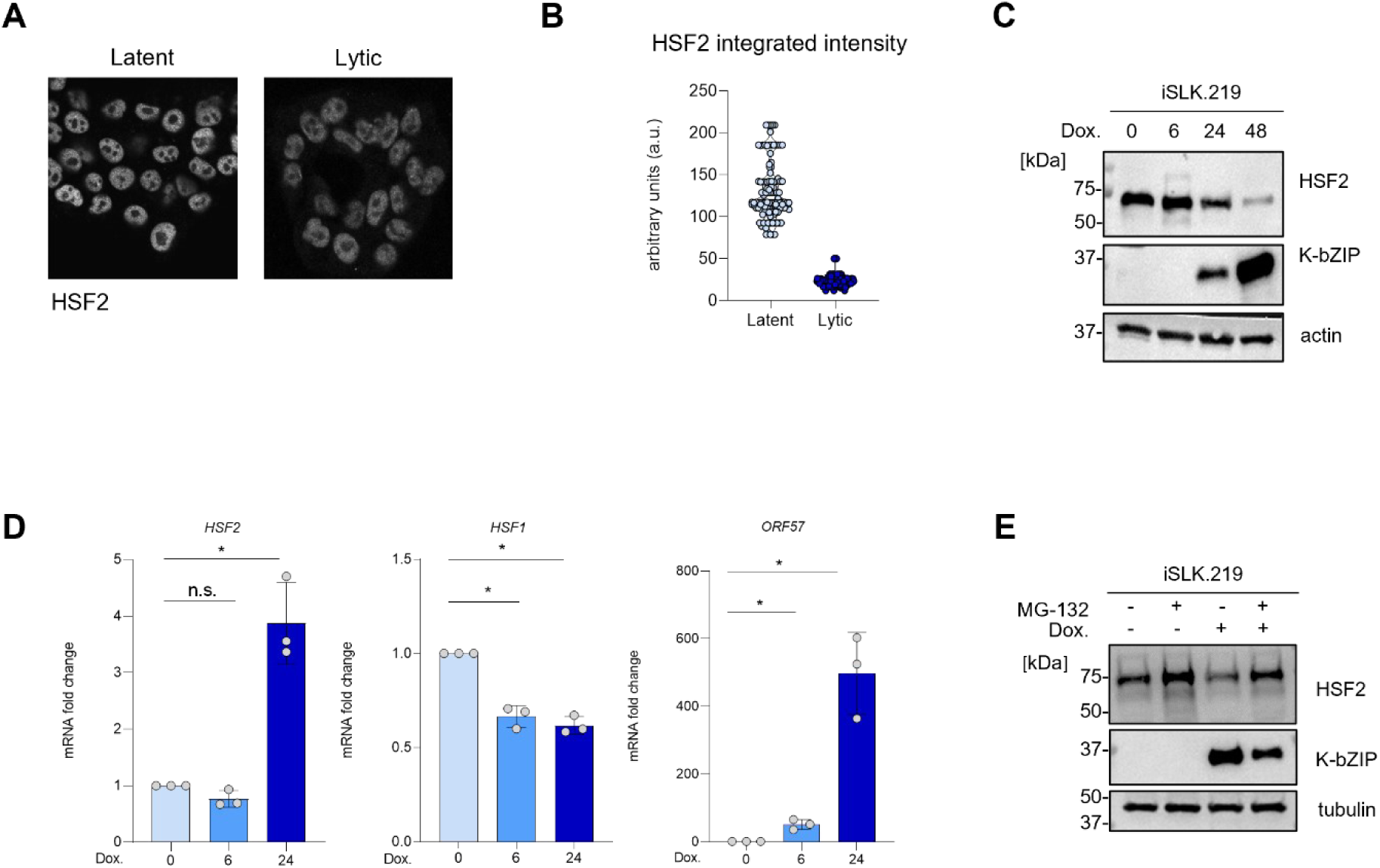
HSF2 is degraded during KSHV lytic reactivation. (A) Confocal representative images and (B) quantification of the HSF2 nuclear integrated HSF2 intensity in n>100 cells in iSLK.219 either latent or induced to the lytic cycle with doxycycline for 24h (indicated as lytic) and stained with an antibody against HSF2. (C) Representatives immunoblot analysis of iSLK.219 cells treated with doxycycline (Dox) for the indicated hours. Levels of HSF2 are shown, K-bZIP and actin are used as reactivation marker and loading control, respectively. Molecular weight in kDa is shown on the left. (D) Trancript analysis of *HSF2*, *HSF1* and viral *ORF57* genes in iSLK.219 cells reactivated with doxycycline (Dox) for the indicated hours. *Actin* was used as internal control. Bars represent the average and error bars the SD across three independent experiments, data points are indicated as grey circles. (E) Representatives immunoblot analysis of iSLK.219 cells latent or induced with Doxycycline for 22 h and were indicated treated with the MG132 proteasome inhibitor (+) or DMSO solvent control (-) for 3 h prior to lysis. HSF2 protein levels are shown, K-bZIP and *tubulin* were used as lytic marker and loading control respectively. Molecular weight in kDa is shown on the left.

During the lytic reactivation, cellular transcripts are degraded as a result of a process called viral-mediated host shut-off (48,49). Earlier studies have estimated that approximately 80% of host transcripts undergo rapid turnover due to KSHV-mediated host shut-off. The viral exonuclease coded by the ORF37 gene and expressed during KSHV lytic phase is responsible for this process (49). The host shut-off is necessary to ensure immune evasion and the preferential usage of the cellular translation machinery to produce viral proteins. Only a minor group of cellular transcripts escape the host shut-off, but the full list of escapees and the mechanisms behind this phenomenon are still incomplete (50,51). We reasoned that the reduction of HSF2 protein levels documented in lytic cells could result from the increased transcript turnover due to the ongoing host shut-off. To verify this result, we measured HSF2 mRNA levels over a 24-h time-course in iSLK.219 cells induced into the lytic cycle by doxycycline treatment. To our surprise, no reduction in the levels of the HSF2 mRNA was detected (Fig. 5D). When measuring HSF2 transcript levels, we found a significant increase in expression 24 h following lytic reactivation, whereas HSF1 levels were downregulated during the analyzed time-course. This result indicated that HSF2 (but not HSF1) may belong to the small group of host transcripts that escape KSHV-induced host shut-off.

Since HSF2 protein downregulation was not due to a reduced transcript turnover, we decided to investigate whether HSF2 downregulation was the result of proteasomal degradation. Interestingly, previous studies have shown that the levels of HSF2 protein can be regulated through the ubiquitin-proteasome pathway. For example during acute stress responses as well as in mitotically arrested cells, HSF2 is ubiquitinated and degraded by the proteasome, in a process mediated by the ubiquitin E3 ligase anaphase promoter complex APC/C (52) (53). To investigate whether HSF2 was subjected to proteasomal degradation during KSHV lytic phase, we treated iSLK.219 cells with the proteasome inhibitor MG132 for 3 h in both latent and lytic phase (treated with doxycycline for 24 h) and measured the levels of HSF2 by immunoblot. Results show that in lytic cells, inhibition of the proteasome activity rescued the levels of HSF2, hence implying that during the lytic replication cycle HSF2 protein levels are diminished as a result of proteasomal degradation.

Taken together, these results suggest that HSF2 levels are subjected to multiple regulatory modalities during lytic KSHV infection: while HSF2 transcript levels are significantly higher possibly due to increased transcription and/or host shut-off escape, the HSF2 protein levels are diminished because of proteasomal degradation.

### HSF2 supports EBV gene expression

Several cellular factors modulate the reactivation of both gamma herpesviruses, EBV and KSHV, underscoring the existence of conserved mechanisms of regulation. Examples include, but are not restricted to, the Barrier-to-autointegration factor 1 (BAF1), Polo-like kinase 1 (PLK1) Aurora Kinase and the mitochondrial protein TBRG4 (54–56). Based on our findings uncovering the positive role of HSF2 in KSHV reactivation, we asked whether EBV would also be subjected to a similar regulatory mechanism. It has been demonstrated that EBV displays a spontaneous lytic cycle in epithelial cells (57–59). Due to its recently discovered role as causal agent of a molecular subtype of gastric cancer (7), we decided to test the effect of HSF2 in EBV life cycle using from EBV-associated gastric cancer primary cells derived from patient-derived-xenografts (PDXs). These cells have been derived from EBVaGC patients, transplanted as xenografts in mice before culturing them *in vitro* (60). We treated these cells with siRNA targeting HSF2 or a control, scr siRNA for 72 h and assessed EBV viral gene expression by RTqPCR. All analyzed viral genes were downregulated including latent (*EBNA1*), the early lytic (*BZRLF1* and *BRLF1*), and the late lytic (*BDLF1*) (Fig. 6A).

**Figure 6.**
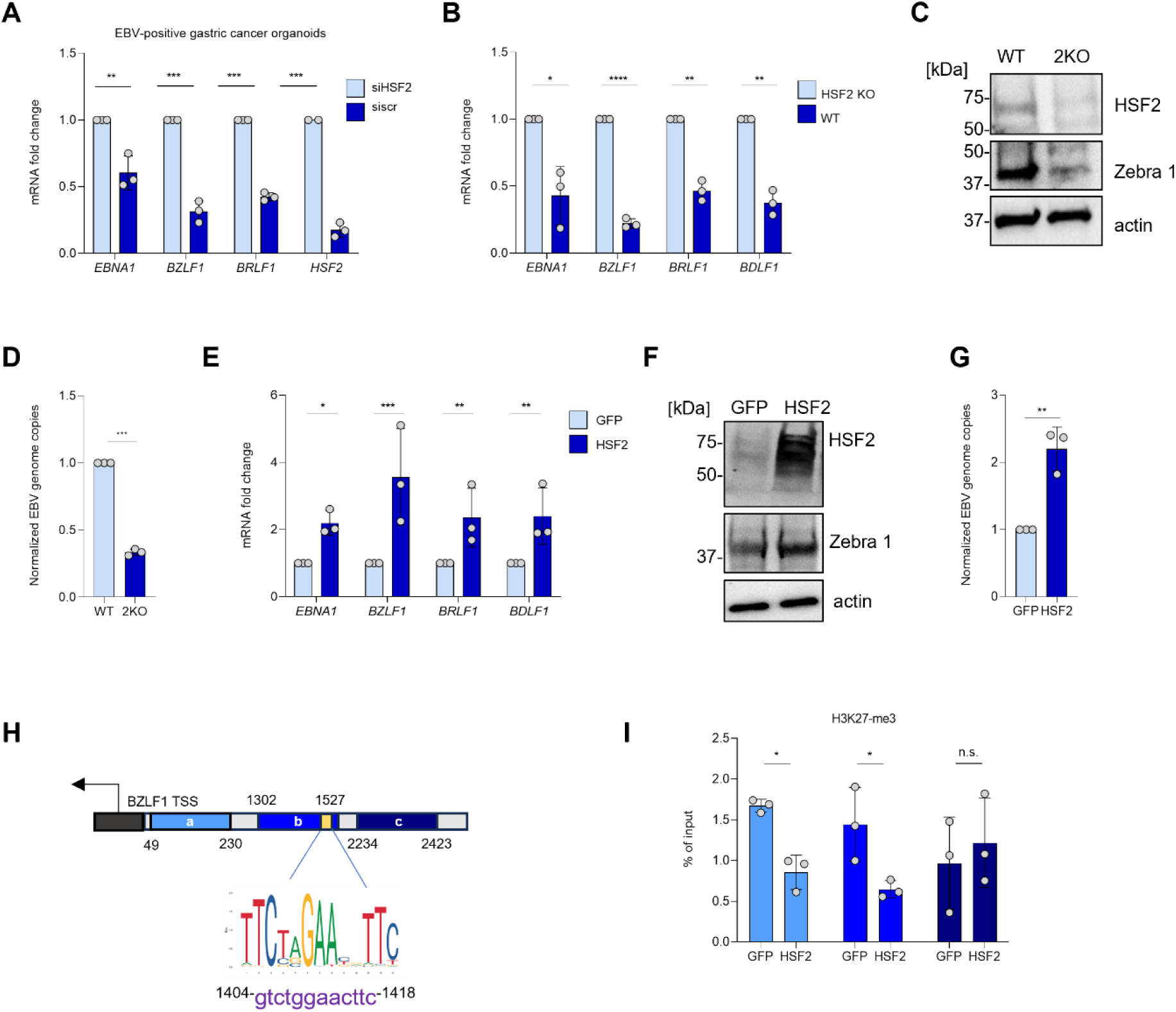
HSF2 is a positive regulator of EBV spontaneous lytic cycle in gastric epithelial cells. (A) patient-derived EBV-positive gastric adenocarcinoma cells were transfected with control, scramble (siscr) siRNA or siRNA targeting HSF2 and the levels of the indicated viral transcripts were measured by RT-qPCR. *Actin* was used as internal control. Bars represent the average and error bars the SD across three independent experiments, data points are indicated as grey circles. (B, C, D) Gastric adenocarcinoma AGS cell lines stably infected with EBV and with lentivirus expressing a CRISP/Cas9 and a sgRNA targeting HSF2 (indicated as 2KO). (B) Cells were analyzed for the indicated EBV viral transcripts by RTqPCR. Bars represent the average and error bars the SD across three independent experiments, data points are indicated as grey circles. (C) Cells were analyzed by immunoblot for HSF2 and the viral protein Zebra 1 (encoded by the BZLF1 gene) proteins, *actin* was used as a loading control. Representative images are shown, molecular weight in kDa is shown on the left. (D) DNA was extracted, and the intracellular viral genome copies were measured. Bars represent the average and error bars the SD across three independent experiments, data points are indicated as grey circles. (E, F, G) Gastric adenocarcinoma AGS cell lines stably infected with EBV and with lentiviruses expressing either HSF2 or GFP. (E) Cells were analyzed for the indicated EBV viral transcripts by RTqPCR with *Actin* as internal control. Bars represent the average and error bars the SD across three independent experiments, data points are indicated as grey circles. (F) Cells were analyzed by immunoblot for HSF2 and Zebra protein levels (encoded by the BZLF1 gene) proteins, actin served as a loading control. Representative images are shown, molecular weight in kDa is shown on the left. (G) DNA was extracted, and the intracellular viral genome copies were measured. Bars represent the average and error bars the SD across three independent experiments, data points are indicated as grey circles. (H) Schematics of the EBV-BZLF1 promoter region, transcription start site (TSS and black arrow) and HSF2 binding site (yellow box) are shown. The regions amplified in ChIP-qPCR (blue segments) and their nucleotide positions are labeled. Below the HSE consensus logo and the corresponding nucleotide sequence within the BZLF1 promoter are shown. (I) HSF2 or GFP control overexpressing AGS BX1 cells were used to immunoprecipitated H3K27-me3 followed by RTqPCR for the indicated primers targeting different regions of the BZLF1 promoter. The bars are colored according to the schematic in (H). Bars represent the average and error bars the SD across three independent experiments, data points are indicated as grey circles.

For subsequent analysis, we decided to utilize the AGS model, a gastric adenocarcinoma cell line, ectopically infected with EBV akata strain (AGS BX1) (61). We genetically deleted HSF2 using CRISPR Cas9 and established a stable AGS BX1 HSF2KO cell line (here indicated as 2KO cells). Lower HSF2 levels were demonstrated by immunoblot and when measuring the viral gene expression, we observed a significant downregulation of EBV viral gene expression and reduced levels of the lytic protein Zebra 1 (Fig. 6B, C). Genetic deletion of HSF2 was also accompanied by diminished intracellular viral genome copies (Fig. 6E). To further validate our findings, we assessed the effect of HSF2 overexpression, and, to this aim, we utilized AGS BX1 to generate stable cell lines ectopically expressing either HSF2 or a GFP control lentivirus. In these cells, EBV viral gene expression was significantly higher in HSF2-overexpressing cells when compared to the GFP control (Fig. 6E). Conversely, also the level of the lytic protein Zebra 1 and the intracellular EBV viral copies were higher (Fig. 6F, G). This suggests that HSF2 modulates EBV gene expression in gastric adenocarcinoma cells, which is consistent with our findings in KSHV-infected cells.

We next explored whether HSF2 might influence EBV lytic reactivation by modulating the BZLF1 promoter, the master regulator of EBV lytic cycle. First, we used the JASPAR database to screen the BZLF1 promoter for HSF2-binding sites. Interestingly, we could detect an HSE motif consisting of two consecutive repeats of the pentameric nGAAn consensus sequence upstream of the BZLF1 TSS (Fig. 6H). It has been previously reported that the EBV BZLF1 promoter, like the KSHV ORF50 promoter, displays a bivalent chromatin structure that allows for quick lytic reactivation upon appropriate trigger (62). These observations led us to ask whether the local epigenetic landscape at the BZLF1 promoter would be different in AGS BX1 cells overexpressing HSF2 than in parental cells overexpressing control GFP., We performed ChIP-PCR to assess H3K27-me3 distribution in one region proximal to the BZLF1 TSS and two more distal regions, with one of them harboring the putative HSF2-binding motif (Fig. 6H). We observed significantly lower levels of the H3K27-me3 repressive chromatin mark in the proximal region and in the distal region harboring the putative HSE within the BZLF1 promoter (Fig. 6I).

In conclusion, these results indicated that HSF2 regulates EBV gene expression in gastric cancer cells and remodels the local chromatin landscape towards a more transcriptionally permissive status at the BZLF1 promoter. Altogether, these findings are indicative of a conserved role for HSF2 acting as a positive modulator of oncogenic gamma-herpesvirus gene expression.

## Discussion

The heat shock factor family of transcription factors is composed of seven HSF members, of which only HSF1 and partially HSF2 are involved in maintaining protein homeostasis under external stress (63). HSF1 is the master regulator of the heat shock response, and it interacts with HSF2 to drive expression of specific genes (24,25,63). Beyond mitigating damage caused by proteotoxic stress, HSF family members are also required for cell differentiation and development of specific organs and their malfunction leads to diverse pathological conditions (21). While the functions of HSF1 have been extensively studied, HSF2 has remained less explored. In this study, we interrogated the role of HSF2 in gamma-herpesvirus reactivation. We show that HSF2 maintains a permissive transcriptional environment in the chromatin that facilitates lytic gene expression in both oncogenic human gamma-herpesviruses, EBV and KSHV. HSF2 modulates viral gene expression in physiologically relevant models of viral-mediated oncogenesis, including primary KLEC and EBV-positive, patient-derived gastric cancer organoids. In addition, HSF2 can affect viral gene expression already in latency when it binds the promoter of the ORF50 gene.

The tight interconnection and cross-regulation between the two HSFs, has provoked us to ask whether HSF1 would contribute to the molecular processes regulated by HSF2. It is worth noting that the interdependence between HSF2 and HSF1 has been examined especially in HSF2-focused studies, while HSF1 research tends to ignore the potential impact of HSF2. This implies that the functions and roles of HSF2 in several cellular and viral processes could be underestimated, warranting further investigations. In this study we found that HSF2 regulates the ORF50 expression independently of HSF1. This is consistent with the features of the HSE located at the ORF50 promoter. The variation in the affinity of HSF1 or HSF2 to the DNA depends on several factors including the HSE architecture, *i.e.* length and orientation of the pentameric nGAAn sequence. *In vitro* and *in vivo* studies have elegantly shown a predilection by HSF2 for shorter HSEs, with only two or three repeats of the pentameric motif, whereas HSF1 preferentially binds cooperatively to longer HSEs (30,64). Accordingly, the ORF50 promoter, harboring a triad of the pentameric nucleotide consensus sequence, represents a better target for HSF2 rather than for HSF1 binding. Similarly, we detected a short HSE motif on the BZLF1 promoter, master regulator of EBV reactivation, suggestive of a possible HSF2-binding site and a conserved mechanism of regulation.

Cells ectopically expressing HSF2 displayed a change in the epigenetic landscape at the ORF50 and BZLF1 promoter regions, in which the levels of the repressive H3K27-me3 chromatin mark were significantly reduced. It has been previously demonstrated that the ORF50 and BZLF1 promoters bear epigenetic features of bivalent chromatin, exhibiting both activating and repressive marks (11,45,62). This poised chromatin state, first identified in lineage-specific promoters of embryonic stem cells (ESCs), was subsequently found at promoters of genes important for somatic development (reviewed in (47,65)). Bivalent chromatin state is also retained in germ cells from fetal stages through meiosis and gametogenesis (66). Interestingly, HSF2 expression is enriched in testes, where it displays a nuclear localization and its genomic depletion leads to defects in spermatogenesis in mice (67–71). Although no direct evidence is available to date, it is possible that HSF2 would be a mediator of chromatin remodeling by regulating the levels of H3K27-me3 at target poised promoters, especially during differentiation processes.

It was previously shown that HSF2 binds to the HSP70 promoter (22,53), and this binding is thought to prevent chromatin condensation, keeping this genomic locus accessible for rapid activation in case of acute stress (72). The HSF2-mediated priming of the HSP70 promoter is important during mitosis when the transcription is dramatically repressed and the chromatin tightly condensed (53). Intriguingly, a few genes escape this silencing and one of them is the gene encoding HSP70. During mitosis, HSF2 is released from the HSP70 promoter, making it accessible to its more potent transactivator HSF1 (22,53). The role of HSF2 at the ORF50 promoter in latent cells shows similarities to its function at the HSP70 promoter. In both cases, HSF2 binds to target promoter regions when the genomic chromatin is repressed. Furthermore, upon lytic reactivation HSF2 is degraded, making the ORF50 promoter accessible to RTA, its main transactivator. This similarity suggests that HSF2 may play a conserved role in facilitating promoter accessibility, preparing the ORF50 promoter for activation by RTA, analogously to how it primes the HSP70 promoter for HSF1.

Our results demonstrated that when the lytic cycle is triggered in KSHV-infected cells, HSF2 is released from the ORF50 promoter and undergoes proteasomal degradation. Similarly, following its dissociation from the HSP70 promoter, HSF2 is subjected to ubiquitin-mediated proteasomal degradation. In mitotic cells, this process is catalyzed the anaphase promoting complex APC/C that was found to be the specific E3 ubiquitin ligase in this pathway (52). Other herpesviruses, like HCMV, induce the disruption of the APC/C, a process required for efficient viral replication (73). In contrast, KSHV lytic cycle does not interfere with the APC/C activity and hence, this E3 ligase may contribute to HSF2 degradation in this setting (74). However, other KSHV lytic proteins, including RTA, feature E3 ubiquitin-ligase activity and could as well target HSF2 for proteasomal degradation, specifically during the lytic cycle (75). Thus, the molecular machinery involved in the proteasomal degradation of HSF2 during lytic reactivation remains to be established.

Recent transcriptomic profiling experiments have reproducibly shown that in tumors of epithelial and endothelial origin, like EBVaGC, NPC and KS, the viral gene expression landscape includes both latent genes and a scattered repertoire of lytic genes (7,76–78). This unusual expression pattern cannot be ascribed to any of the canonical viral expression programs, and it has been referred to as permissive latency, a “leakage” of lytic genes within the latent program, and more recently it has been defined as a “mixed” expression program (8,9,77,78). These observations are further reinforced by the independent detection of multiple viral lytic proteins in tumor biopsies of KS lesions. Lytic proteins like K8.1 and K15 have been detected together with a robust expression of the lytic viral tegument gene ORF75 (37,77,79,80). Interestingly, the lytic expression pattern found in KS lesions did not correlate with the levels of plasma viremia, suggesting that even in the lesions with higher levels of lytic gene expression, KSHV infection was not productive (80). This implies, at least in these tumors, the presence of complex patterns of viral gene expression different from the classically and mutually exclusive latent vs. lytic programs. Hence, we propose that HSF2 by promoting ORF50 gene expression, would contribute to the “leakage” of downstream lytic transcripts in latently infected cells within these tumors. In this context, HSF2 could represent a pro-oncogenic host factor supporting the expression of viral lytic oncogenes outside of the traditional productive lytic cycle.

Collectively, our findings provide evidence of a novel role for HSF2 as a previously underestimated, common player in EBV and KSHV gene regulation. Specifically, we show that HSF2 induces a more permissive chromatin environment at the RTA and BZLF1 promoters and the expression of lytic genes prior to the initiation of the lytic cycle. These results add HSF2 to the complex host molecular network which controls gamma herpesviral latent-to-lytic switch, a process strongly connected to oncogenesis. Furthermore, this study provides new clues on the function of HSF2 at poised gene promoters, which could advance our understanding of its role in cellular differentiation processes.

## Materials and Methods

### Cell lines

HEK293FT (Thermo Fisher Scientific, R70007; RRID:CVCL_6911) and U2OS cells WT and *Hsf2*^−/−^ (2KO) (22,81) cells were maintained in Dulbecco’s modified Eagle media (DMEM) supplemented with 10% heat-inactivated FBS (GIBCO) 1% L-glutamate and 1% Penicillin/streptomycin.

iSLK.219 (38), were maintained in DMEM media supplemented with 10% heat-inactivated FBS (GIBCO) 1% l-glutamate and 1% penicillin/streptomycin in the presence of puromycin (0.01 mg/mL) (A11138-03, GIBCO), hygromycin B (1.2 mg/mL) (10687010, Invitrogen) and G418 (0.8 mg/mL) (G8168, Sigma Aldrich). When plated for experiments or virus production antibiotics were not supplied to the media.

Primary human dermal lymphatic cells (C-12216, PromoCell) were grown in endothelial basal media (CC-3162, Lonza) supplemented with EGM-2 MV Microvascular Endothelial SingleQuots (CC-4176).

AGS BX1 were maintained in Roswell Park Memorial Institute (RPMI) 1640 (R5886-500, Sigma-Aldrich) media supplemented with 10% heat-inactivated FBS (GIBCO) 1% L-glutamate and 1% penicillin/streptomycin in the presence of puromycin (5 μg/mL) and G418 (0.3 mg/mL) for the antibiotic selection. AGS BX1 cell lines were a kind gift from Maria Masucci (Karolinska Institutet, Sweden).

Gastric cancer primary cells were obtained from Silvia Giordano (University of Turin) and characterized as reported in (60). Cells were grown in ISCOVE media (GIBCO) supplemented with 10%FBS and 1% penicillin/streptomycin on collagen (rat collagen type 1 solution, Sigma Aldrich) coated plates (60).

All cells were maintained under standard conditions (37C, 5% CO_2_ and in a humidified environment)

### KSHV reactivation and virus production

iSLK.219 cells were seeded to reach a 70-80% confluency and the next day the lytic cycle was induced with doxycycline (1 ug/mL). To demonstrate the efficiency of reactivation images were taken with ZOE Fluorescent Cell Imager. For virus production, media was harvested 96 hours post-reactivation and centrifuged at 1000 RCF at 4°C for 30 minutes. rKSHV.219 particles then were concentrated with PEG-it (LV825A-1, System Biosciences) according to the manufacturer’s instructions.

### Lentiviral production and lentiviral transduction

HEK293FT cells were plated on a T75 flask to reach 60-70% confluency the next day. One day after plating, cells were transfected with the pLP1, pLP2 and VSVg plasmids (Addgene) and with the lentiviral plasmid carrying the gene of interest using lipofectamine 3000 (L3000008, Invitrogen) according to the manufacturer’s instructions. Media was changed the next day and lentivirus-containing media was harvested at 72h and 96h after transfection. Lentivirus particles were concentrated with PEG-it (LV825A-1, System Biosciences) according to the manufacturer’s instructions. Lentiviral expression vectors were designed and purchased from VectorBuilder Inc. HSF2 knockout: hHSF2[gRNA#1151]; HSF2 overexpression and control: hHSF2[NM_004506.3]; GFP: pLV[Exp-Puro-EF1A>EGFP (28).

Lentiviral transduction was performed on 60-70% cell layers by adding the lentivirus to fresh growth media in the presence of 8 ug/mL of polybrene followed by spinoculation at 450g for 30 minutes at RT.

### Generation of stable cell lines

1×10^5^ AGS BX1 cells were plated in a 6-well plate and transduced with the indicated lentiviruses. Cells were maintained as a bulk culture and puromycin selection (5 μg/mL) was initiated 48 h after transduction.

U2OS rKSHV.219 cells were generated by infecting U2OS cells with rKSHV.219 virus by spinoculation at 450g, for 30 min at RT. One day post-infection puromycin selection (5 μg/mL) was initiated. Klec were generated as reported in (37).

Cells were maintained in selection antibiotics-containing media. Selection antibiotics were not added when cells were plated for experiments

### siRNA transfection

U2OS rKSHV.219, iSLK.219, KLEC cells were transfected with either 10 pmol/mL siRNA ON-target plus smart pool targeting HSF2 (Dharmacon, L-011874-00-0020), HSF1 (Dharmacon, l-012109-02-0005) or control, scrambled siRNA (Dharmacon, D-001810-10) in presence of the lipofectamine RNAiMAX transfection reagent (13778-075, Invitrogen) according to the instructions provided by the manufacturer. The siRNA used is a mixture of 4 different siRNA provided as a single reagent, thus ensuring both higher potency and specificity than an individual siRNA. The efficacy of HSF2 and HSF1 silencing was confirmed by immunoblotting.

### Luciferase reporter assay

3×10^4^HEK-293FT cells/well were seeded in a 96-well white plate. The next day cells were co-transfected in triplicate with 50 ng of the reporter plasmids, or the corresponding control vector and either 100 ng of HSF2 or control GFP expressing plasmids and where indicated, pFuW-Myc-ORF50 plasmid (82) in the presence of Fugene HD (E2311, Promega) according to the manufacturer’s instructions. 48 hours after transfection cells were lysed in 50 µl/well of ONE-GloLuciferase Assay Reagent (E6110, Promega). The luciferase activity was measured by total counts acquired using the HIDEX sense 425-301i plate reader and software (Hidex).

The following reporter plasmids were used in this study: pGL2-basic (Promega); pGL2-ORF50 pGL3-basic (Promega); pGL3-7XTR; pGL3-OriLyt; pGL3-ORF45 and pGL3-K8.1 (provided by T.F. Schulz, Hannover Medical School, Germany).

For each experimental setting, three independent experiments were performed in triplicates. The transfection efficiency, for each condition, was assessed with immunoblotting.

### Quantification of intracellular viral genome copies

DNA was isolated using QIAamp DNA Micro Kit (56304, Qiagen) and analyzed by qPCR using primers for genomic actin (5’-AGAAAATCTGGCACCACACC-3’; 5’-AACGGCAGAAGAGAGAACCA-3’) and genomic EBV copies were detected by amplification of a portion of the gp85 region (5’-TGTGGATGGGTTTCTTGGGC-3’; 5’-TGGTCAGCAGCAATAGTGAAGC-3’).

### Inhibitor treatment

0.5×10^5^ iSLK.219 cells/well were seeded in a 24-well plate. On the next day, cells were reactivated with Doxycycline. 22 hours post reactivation cells were either treated with the protease inhibitor MG-132 (2194S, Cell Signaling Technology) used with a concentration of 20 μM or an equal volume of DMSO used as negative control. 24 hours post reactivation cells were harvested and stored at -80°C for further analysis.

### qRT-PCR

RNA was isolated with RNeasy Mini Kit (74136, QIAGEN) according to the manufacturer’s instructions. The amount of RNA was quantified using a NanoDrop 2000 spectrophotometer (ThermoFisher Scientific), and for each sample, 900 ng of RNA were reverse transcribed using the iScript cDNA Synthesis Kit (#1708891, Bio-Rad). SensiFAST SYBR Hi-ROX kit (BIO-92020, Meridian Bioline) was used in the qPCR reactions, which were performed using a QuantStudio 3 Real-Time PCR system (Applied Biosystems, Thermo Fisher Scientific). Actin or 18S were used as housekeeping genes for the normalization of all other gene expression. Samples were run in triplicates and each experiment was done at three independent times. The primer sequences are listed in Table 1.

### ChIP-PCR

SimpleChip Enzymatic Chromatin IP Kit (Magnetic Beads) (#91820:9003, Cell Signaling Technology) was used according to the manufacturer’s instructions. For each immunoprecipitation (IP), the chromatin from 6x 10^5^ iSLK.219 or AGS BX.1 cells was crosslinked with 1% formaldehyde.

For the identification of HSF2 association on specific regions of the KSHV ORF50 promoter 2μL of antibodies against HSF2 (SFI58)(22) and as control normal Rabbit IgG (#2729, Cell Signaling Technology) were used. For the quantification of histone markers tri-methyl-histone H3(K4) (#9751, Cell Signaling technology), tri-methyl-histone H3(K27) (#9733T, Cell Signaling technology) and normal Rabbit IgG (#2729, Cell Signaling Technology) were used at the concentration of 1:150 as per manufacturer’s recommendation.

The experiments were performed at least three independent times. Isolated and purified DNA was amplified by quantitative real-time PCR using the indicated primer pairs listed in Table 2

### Immunoblotting

Cells were lysed in 3x Laemmli lysis buffer (30% glycerol, 187.5 mM SDS, 3% Tris-HCl, 0.015% bromophenol blue, 3% β-mercaptoethanol), and boiled for 5 minutes at 95 °C and then resolved on 7.5% or 4-20% Mini-PROTEAN TGX Stain-Free Precast Gels (Bio-Rad) and transferred onto an Amersham Protran 0.45 nitrocellulose membrane (Cytiva). The membranes were blocked with 5% milk-PBS-Tween20 to detect the proteins of interest. The following primary antibodies were diluted in 5% milk-PBST and used for immunoblotting with a 1:1000 dilution for the anti-HSF2 (HPA031455, Sigma-Aldrich) and anti-HSF1 (SMC-118C, StressMarq Biosciences), 1:500 dilution for anti-K8.1 (sc-65446, SantaCruz); anti-K-bZIP (sc-69797, SantaCruz); 1:250 anti-LANA (AB4103, abcam); anti-actin (A4700, Sigma-Aldrich); anti-alpha-tubulin (12G10, DSHB) and incubated overnight at 4°C. After three rounds of washing in PBST the following Horseradish peroxidase-conjugated secondary antibodies were diluted in 5% milk-PBST and used with at a 1:10000 dilution anti-rabbit (W4018, Promega), 1:5000 anti-mouse (W402B, Promega), 1:3000 anti-Rat (ab97057, abcam) and incubated at room temperature for 1 hour. After 3 rounds of washing in PBST luminescent signal was revealed with Super Signal West Pico PLUS Chemiluminescent Substrate (34580, Thermo Scientific) using the chemiluminescent programme of detection on the iBrightCL1000. Experiments were done at least two independent times; representative experiments are shown. Equal loading of the independently generated blots was ensured by Ponceau-S staining.

### Immunofluorescence

iSLK.219 cells plated on coverslips were either treated with Doxycycline or left untreated. 24-hour post reactivation cells were fixed with 4% paraformaldehyde for 10 min at RT and washed three times with PBS. Cells were then permeabilized and stained with DAPI (1 ug/mL) dissolved in 0.5% Triton X-100 and 3 mM EDTA in PBS in the dark, at room temperature, for 10 min. The cells were then washed three times with PBS and blocked with 1% BSA in TBST for 1 h at RT and incubated with primary antibodies overnight at 4°C. The primary antibodies were diluted in 1% BSA in TBST: 1:100 anti-HSF2 (HPA031455, Sigma-Aldrich). After primary antibody incubation, cells were washed with PBS incubated with secondary antibodies for 1 hour at RT. The following secondary antibodies were diluted in 1% BSA-TBST; 1:500 anti-rabbit Alexa Fluor 547(A-21244, Invitrogen). Next, the cells were washed with PBS and with MQ-water and then mounted with VECTASHIELD mounting medium (H-100, Vector Laboratories). Coverslips were imaged with Zeiss LSM 880 microscope using a 40X objective. Acquired images are shown as maximum intensity projections. The integrated nuclear intensity was used to compare the HSF2 levels between the latent and lytic-infected cells.

For viral titre quantification, U2OS cells were plated on a 96 clear bottom black well-plate (655090, Grainer) infected with serial dilution of supernatant from KLEC treated as indicated for 24h. Cell were then fixed and stained with DAPI. The primary antibodies were diluted in 1% BSA in TBST: 1:200 anti-LANA (AB4103, abcam) and the secondary antibody used was anti-rat Alexa Fluor 647 (A-21247, Invitrogen) diluted 1:500. Stained cells were imaged with an HT microscope using a 10X objective. Cells positive for LANA were identified by subtracting the background intensity from uninfected cells (stained with LANA antibody) and expressed as percentage of LANA positive cells. The number of LANA-positive cells were quantified over the total amount of nuclei (number of nuclei) using Cell Profiler (http://cellprofiler.org)

### Statistical analysis

Data were analyzed with Prism GraphPad 8. Unless differently stated in the legend, the graphs show the single values of each biological replicate, the mean and the error bars indicate the SD across the biological replicates. Ordinary one-way ANOVA followed by Dunnett correction for multiple comparisons or two-tailed paired *t* test were performed to assess the statistical significance of the differences between samples. *: p<0.05; **: p<0.01; ***: p<0.001; n.s.:non-significant.

## Acknowledgement

We thank the members of the Sistonen research group for critical discussions. Prof. Maria Masucci, Prof. T.F. Schulz and Prof. Päivi M. Ojala are acknowledged for providing plasmids and cell lines. The staff of the facilities of the Turku Screening Unit, a member of the Biocentre Finland Drug Discovery and Chemical Biology Network and EU-Openscreen ERIC and Turku Bioscience Center are acknowledged for technical support.

The study was supported by Finnish Society of Sciences and Letters (SGr), Mary and Georg Ehrnrooth Foundation (SGr) and K. Albin Johansson Foundation (SGr), Sigrid Jusélius Foundation (LS), Cancer Foundation Finland (LS), The Medical Foundation Liv och Hälsa (LS). LC, LS, SGr were supported by the Research Council of Finland; grant number 355708 (SGr, LC), 355596 (LS), SGr was also supported by the Finnish Cultural Foundation. SC and SGi were supported by the Italian association for Cancer research (AIRC) IG 27531.

